# Genomic selection analyses reveal tradeoff between chestnut blight tolerance and genome inheritance from American chestnut (*Castanea dentata*) in (*C. dentata* x *C. mollissima*) x *C. dentata* backcross populations

**DOI:** 10.1101/690693

**Authors:** Jared W. Westbrook, Qian Zhang, Mihir K. Mandal, Eric V. Jenkins, Laura E. Barth, Jerry W. Jenkins, Jane Grimwood, Jeremy Schmutz, Jason A. Holliday

## Abstract

American chestnut was once a foundation species of eastern North American forests, but was rendered functionally extinct in the early 20th century by an exotic fungal blight (*Cryphonectria parasitica*). Over the past 30 years, The American Chestnut Foundation (TACF) has pursued backcross breeding to generate hybrids that combine the timber-type form of American chestnut with the blight tolerance of Chinese chestnut. The backcross strategy has been implemented based on the hypothesis that blight tolerance is conferred by few major effect alleles. We tested this hypothesis by developing genomic prediction models for five presence/absence blight phenotypes of 1,230 BC_3_F_2_ selection candidates and average canker severity of their BC_3_F_3_ progeny. We also genotyped pure Chinese and American chestnut reference panels to estimate the proportion of BC_3_F_2_ genomes inherited from parent species. We found that genomic prediction from a method that assumes an infinitesimal model of inheritance (HBLUP) has a similar predictive ability to a method that tends to perform well for traits controlled by major genes (Bayes C). Furthermore, the proportion of BC_3_F_2_ trees’ genomes inherited from American chestnut was negatively correlated with the blight tolerance of BC_3_F_2_ trees and their progeny. On average, selected BC_3_F_2_ trees inherited 83% of their genome from American chestnut and have blight-tolerance that is intermediate between F_1_ hybrids and American chestnut. Results suggest polygenic rather than major gene inheritance for blight tolerance. The blight-tolerance of restoration populations will be enhanced by advancing additional sources of blight-tolerance through fewer backcross generations and by potentially by breeding with transgenic blight-tolerant trees.

## Introduction

Efforts to restore the American chestnut (*Castanea dentata*) have been ongoing for nearly 100 years. The chestnut blight fungus (*Cryphonectria parasitica*), first introduced into North America from Asia in the early 1900s, killed approximately 4.2 billion *Castanea dentata* stems from northern Mississippi to coastal Maine by the 1950s (Little, 1977; Gravat, 1949; Hepting, 1974; Newhouse, 1990). The extirpation of *C. dentata* reduced wildlife carrying capacity and altered nutrient cycling in forests throughout its native range (Ellison et al., 2005; Dalgleish et al., 2012). Today, an estimated 431 million American chestnuts stems survive as seedlings and collar sprouts, but their stems rarely flower and almost never produce viable seed before being re-infected with the blight (Dalgliesh et al., 2016). Publicly funded breeding programs, initiated in the 1920s by the U.S. Department of Agriculture and the Brooklyn Botanical Garden, hybridized *C. dentata* with Asian *Castanea* species that are tolerant of chestnut blight (Burnham et al., 1986; Anagnostakis, 2012). However, these F_1_ hybrids were not sufficiently competitive in the mixed hardwood forests typical of the historical *C. dentata* range (Schlarbaum et al., 1998), and these early chestnut breeding programs were largely discontinued by the 1960s (Jaynes, 1978).

In 1983, The American Chestnut Foundation (TACF) was founded and backcross breeding was proposed to generate hybrids that combined the blight-tolerance of Chinese chestnut (*Castanea mollissima*) with the timber-type form of American chestnut (Burnham, 1981; Burnham et al., 1986; Burnham, 1988). Backcrossing *C. mollissima* x *C. dentata* hybrids to *C. dentata* over three generations was expected to generate BC_3_ hybrids that inherited an average of 15/16^ths^ (93.75%) of their genome from *C. dentata*. The BC_3_ trees were intercrossed to generate BC_3_F_2_ populations from which a subset of trees were predicted to be homozygous for blight-tolerance alleles from *C. mollissima*. Large quantities of blight-tolerant BC_3_F_3_ seed for restoration would then be generated through open-pollination among the selected homozygous blight-tolerant BC_3_F_2_ trees.

The backcross method was initially implemented based on two hypotheses. First, alleles for blight-tolerance segregate at a few loci with incomplete dominance. Second, trees that are heterozygous for blight-tolerance at all loci can be reliably selected in each backcross generation. Incomplete dominance of blight-tolerance was surmised from the observation that F_1_ hybrids develop blight cankers that are intermediate in size and severity between *C. mollissima* and *C. dentata* (Graves 1950). Burnham et al. (1986) hypothesized that blight-tolerance segregates at two loci based on observations of Clapper (1952) that F_1_ hybrids backcrossed to *C. mollissima* segregate at a ratio of three small cankered trees (blight-tolerant) to one large cankered tree (susceptible). Later, Kubisiak et al. (1997; 2013) found that three QTLs on three linkage groups (B, F, G) explained 40% of the variation in canker severity in a full-sib (*C. dentata* x *C. mollissima*) x (*C. dentata* x *C. mollissima*) F_2_ family.

TACF began backcross breeding in 1989 by pollinating two (*C. dentata* x *C. mollissima)* x *C. dentata* BC_1_ hybrids (the ‘Clapper’ and ‘Graves’ trees) with *C. dentata* pollen from multiple trees in southwest Virginia (Hebard, 2006; Steiner et al., 2017). These BC_1_ trees were chosen as sources of blight-tolerance to reduce the number of additional generations of breeding and selection required to reach the BC_3_F_3_ generation. The ‘Clapper’ and ‘Graves’ trees have different *C. mollissima* grandparents (Clapper, 1963; Hebard, 2006), and were bred as distinct sources of resistance based on the possibility that blight-tolerance would segregate at different loci among the progeny of these trees. Phenotypic selection was performed in the BC_2_ and BC_3_ generations at TACF’s Research Farms in Meadowview, Virginia, by artificially inoculating stems with *C. parasitica* and selecting trees with subjective canker severity ratings that were indistinguishable from F_1_ hybrids (Steiner et al., 2017). Additional selection was made for leaf and twig characteristics that resembled those of *C. dentata* (Hebard, 1994; Diskin et al., 2006). Citizen scientists affiliated with TACF have subsequently pollinated wild-type trees ranging from Alabama to Maine with pollen from selected BC_2_ and BC_3_ trees from the Meadowview breeding program to increase the genetic diversity and adaptive capacity of backcross populations (Westbrook, 2018; Fig. 1).

**Figure 1:**
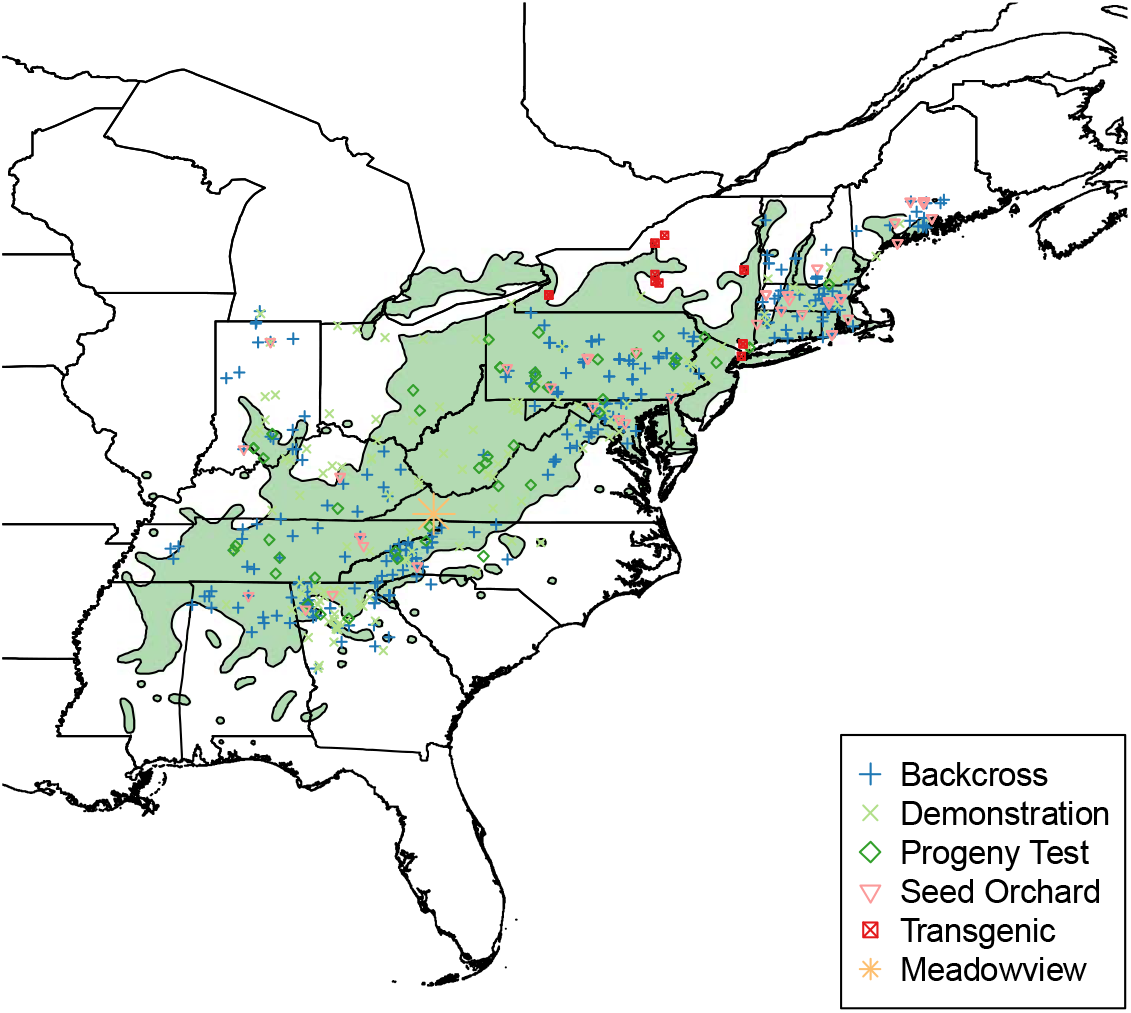
Map of The American Chestnut Foundation orchard locations across the native range of *Castanea dentata*.

The Meadowview backcross breeding program is now reaching the final stages of selection for blight tolerance. Large segregating BC_3_F_2_ populations have been generated by open-pollination among selected BC_3_ descendants of the ‘Clapper’ and ‘Graves’ trees. Between 2002 and 2018, approximately 36,000 BC_3_F_2_ progeny of 83 ‘Clapper’ BC_3_ selections and 28,000 BC_3_F_2_ progeny of 68 ‘Graves’ BC_3_ selections were planted in two seed orchards (Steiner et al., 2017). Assuming that blight-tolerance segregates at three unlinked loci (Kubisiak et al., 1997), that all BC_3_ selections were heterozygous for *C. mollissima* alleles at these loci, and that 80% of BC_3_F_2_ seeds planted would survive to inoculation, there is a 99% probability of generating nine homozygous blight-tolerant BC_3_F_2_ trees from each backcross line (Hebard, 1994; Hebard, 2002).

Between 60% and 80% of BC_3_F_2_ trees were culled on the basis of significant canker expansion six months after inoculation. Additional culling was performed based on blight phenotypes that take longer to develop, such as the survival of the main inoculated stem and the severity of additional cankers that developed as a result of natural infection by *C. parasitica* (Hebard, 2006). As of 2018, ~3,300 ‘Clapper’, and ~4,300 ‘Graves’ BC_3_F_2_ trees remain. To select the most resistant of these remaining trees, TACF has planted randomized field trials of their open-pollinated progeny. After inoculating these trials with *C. parasitica*, average canker severity of the most blight tolerant BC_3_F_3_ families was intermediate between Chinese chestnut and American chestnut. This finding led Steiner et al. (2017) to hypothesize that blight tolerance segregates at more loci than previously assumed and that phenotypic selection has not been sufficiently accurate to select for the complete set of resistance alleles from *C. mollissima* founders in all backcross lines.

Progeny testing all BC_3_F_2_ mothers is potentially more accurate than selection based on blight phenotypes of individual BC_3_F_2_ trees. However, screening 7600 BC_3_F_2_ remaining mother trees would require planting hundreds of thousands of progeny and waiting many years for all BC_3_F_2_ selection candidates to flower. An alternative approach is genomic selection, which enables simultaneous ranking of a large number of BC_3_F_2_ selection candidates for blight-tolerance including younger trees that have not flowered. In the context of TACF’s breeding program, genomic selection entails genotyping training populations composed of BC_3_F_2_ trees whose progeny have been inoculated with *C. parasitica* or that have been phenotyped for late-developing blight traits. Breeding values for BC_3_F_2_ selection candidates that have not been phenotyped for these traits may be predicted from a blend of pedigree and genomic relationships with trees in the training population using the single step BLUP (or HBLUP) method (Legarra et al. 2009; Misztal et al., 2009; Aguilar et al., 2009). Alternatively, a genome-wide panel of SNP genotypes may be regressed on phenotypes from the training population and breeding values predicted by multiplying marker genotypes by allelic substitution effects (Meuwissen et al., 2001).

In this study, our first aim was to optimize an analytical pipeline for genomic selection for blight tolerance in American chestnut backcross populations. Towards this end, we generated a draft reference genome for *C. dentata* and performed genotyping-by-sequencing on 1,230 BC_3_F_2_ selection candidates from the Meadowview breeding program. We optimized the HBLUP method to predict breeding values for late developing blight phenotypes of BC_3_F_2_ selection candidates and average canker severity of their BC_3_F_3_ progeny. We then summed the breeding values for these traits to create a selection index to compare the blight tolerance of BC_3_F_2_ selection candidates under different selection scenarios.

Our second aim was to test the hypothesis that blight-tolerance from *Castanea mollissima* segregates at a few major effect loci. We tested this hypothesis first by comparing the predictive ability of HBLUP to Bayes C regression. Bayes C, which includes only the largest effect markers in the prediction model, has been found to have greater predictive ability than HBLUP for traits that are controlled few major effect loci, whereas HBLUP and Bayes C have similar predictive ability for polygenic traits (Resende et al., 2012; Chen et al., 2014; Yoshida et al., 2018). We also tested this hypothesis by estimating the correlation between the proportion of BC_3_F_2_ trees’ genomes inherited from *C. dentata* and breeding values for blight tolerance of these trees.

## Materials and Methods

### Phenotyping

#### Phenotyping BC_3_F_3_ progeny

Between 2011 and 2016, 7,173 BC_3_F_3_ progeny from 346 ‘Clapper’ and 198 ‘Graves’ open-pollinated BC_3_F_2_ mothers were evaluated for blight-tolerance. Between 27 and 33 BC_3_F_3_ progeny from each BC_3_F_2_ mother were planted at TACF’s Meadowview Research Farms in a completely randomized design (2011 – 2013 tests) or an alpha-lattice incomplete block design (2014 – 2016 tests) (Patterson & Williams, 1976). In their third growing season, the main stems of BC_3_F_3_ trees were inoculated with the SG2,3 (weakly pathogenic) and Ep155 (highly pathogenic) strains of *C. parasitica* at two stem heights approximately 25 cm apart using the cork borer agar disk method (TACF, 2016). The SG2,3 and Ep155 strains were originally isolated from American chestnut trees in Virginia and Maryland, respectively (M. Double, pers. communication). Inoculation with these two strains increases the range of canker severity phenotypes. However, BC_3_F_3_ family rankings for average canker severity using these two strains have been found to be strongly genetically correlated (r_genetic_ >0.95), suggesting generalized rather than strain-specific mechanisms of host blight tolerance (Steiner et al., 2017; Westbrook & Jarrett, 2018).

Canker lengths and subjective ratings were phenotyped 5 to 6 months after inoculation. Cankers were rated as 1 = minimal expansion beyond initial lesion, 2 = some expansion, but canker partially contained by callus formation, or 3 = canker large, sunken, and sporulating (Fig. A1). The trait ‘canker severity’ was calculated separately for each strain of *C. parasitica* (SG2,3 & Ep155) by scaling the variation in canker lengths and canker ratings to mean 0 and standard deviation 1, and summing the standardized rating and length. The canker severities for each strain of *C. parasitica* were the summed to obtain a single canker severity value for each tree. Canker severity phenotypes were obtained for 48% of the BC_3_F_3_ seeds that were planted and 2 to 40 BC_3_F_3_ progeny (median = 13) were phenotyped per BC_3_F_2_ mother. Canker severity phenotypes of BC_3_F_3_ trees were continuously distributed and there was no difference in the average canker severity in the Clapper and Graves BC_3_F_3_ populations (Fig. A2).

#### Phenotyping BC_3_F_2_ parents

Trees remaining in Meadowview seed orchards that were between 5 and 16 years old were phenotyped for five binary traits hypothesized to be indicative of blight-tolerance or susceptibility. All trees were phenotyped for main stem survival. Trees with a living main stem were then phenotyped for four additional traits on the main stem namely, presence or absence of any canker longer than 15 cm; presence or absence of exposed wood; presence or absence of sporulation of *C. parasitica* conidia from cankers; and presence or absence of sunken cankers. In total, 1134 ‘Clapper’ and 1042 ‘Graves’ BC_3_F_2_ selection candidates for were phenotyped for these traits.

### Marker discovery

#### Generation of a draft reference genome for Castanea dentata

We generated a draft reference genome sequence for the immediate purpose of detecting SNP variants in backcross populations. We sequenced the ‘Ellis1’ clone of *Castanea dentata* by whole genome shotgun sequencing using the PACBIO SEQUEL sequencing platform at the HudsonAlpha Institute in Huntsville, Alabama. A total of 16 cells using chemistry 2.1 were sequenced with a p-read yield of 88.69 Gb (8,327,003 reads), for a total coverage of 98.54x (median read size 7,745 bp). The reads were assembled using MECAT (Xiao et al., 2017) and subsequently polished using ARROW (Chin et al., 2013). This produced 2,959 contigs with an N50 of 4.4 Mb, and a total genome size of 967.1 Mb. Contigs were then collapsed to remove redundant alternative haplotype sequence and screened against bacterial proteins, organelle sequences, and the GenBank non-redundant database to detect and remove contaminants. Version 0.5 of the *C. dentata* genome contains 793.5 Mb of sequence, consisting of 950 contigs with a contig N_50_ of 8.1 Mb.

#### Genotyping-by-sequencing of BC_3_F_2_ trees

Newly expanded leaves were collected from BC_3_F_2_ trees in Meadowview seed orchards in June 2017. The leaf tissue was ground in liquid nitrogen and genomic DNA was extracted using a Qiagen DNeasy Plant Mini kit. The quality and quantity of DNA was checked on a Nanodrop spectrophotometer (ND-100) and 200 ng of DNA from each tree was digested with 1 ul of ApeKI and Illumina-compatible adapters with ApeKI overhangs. Adapters were ligated with 1.6 ul of T4 DNA ligase. Each of the P1 adapters had a variable length (4-8bp) index downstream of the sequencing primer such that it was read immediately preceding the restriction site. The P2 adapter was common across all samples. Following adapter ligation, 18 cycles of PCR were performed to confirm ligation and the fragment size range. The DNA samples were randomly assigned to pools of 50 per lane for trees whose progeny had previously been inoculated with *C. parasitica*, or 96 per lane for trees whose progeny had not been inoculated. Pools were purified with the New England Biolabs Monoch PCR and DNA Clean Up Kit before and after 18 cycles of PCR amplification. Fragments in the range of 250-600 bp were selected on a BluePippin™ instrument (Sage Science, Beverly, MA, USA) and the resulting libraries were visualized on a Bioanalyzer (Agilent 2100 BioAnalyzer). Libraries were then sequenced on an Illumina HiSeq 4000 instrument in 2×150bp paired end mode at the Duke University Center for Genomic and Computational Biology.

Raw reads were filtered for quality, filtered for adapter contamination, and de-multiplexed using STACKS software (Catchen et al., 2013). Filtered reads were then aligned to v. 0.5 of the *C. dentata* reference genome using the Burrows-Wheeler Aligner (BWA) *mem* algorithm, and subsequently converted to BAM format, sorted, and indexed with SAMtools (Li & Durbin 2010; Li et al. 2009). GVCF files for each sample were generated using the GATK HaplotypeCaller algorithm (McKenna et al. 2010; Poplin et al. 2017), and these GVCFs were then merged using the GenotypeGVCFs function to create a candidate polymorphism set. Variants were flagged and removed as low quality if they had the following characteristics: low map quality (MQ < 40); high strand bias (FS > 40); differential map quality between reads supporting the reference and alternative alleles (MQRankSum < −12.5); bias between the reference and alternate alleles in the position of alleles within the reads (ReadPosRankSum < −8.0); and low depth of coverage (DP < 5). The resulting VCF file was filtered to retain only biallelic SNPs with <10% missing data and minor allele frequencies >0.01, leaving 71,507 SNPs. Missing SNP genotypes were imputed with Beagle v 4.1 (Browning & Browning, 2016). A total of 1,230 (865 ‘Clapper’ and 365 ‘Graves’) BC_3_F_2_ individuals were genotyped.

### Genomic prediction and validation

#### Single-step genomic prediction of progeny canker severity

Breeding values for progeny canker severity were obtained for all BC_3_F_2_ mothers that were genotyped and/or whose progeny were phenotyped using the single-step HBLUP method (Legerra 2009; Misztal et al. 2009; Aguilar et al., 2009). This method blends the pedigree and genomic relationship matrix into a single matrix **H** so that phenotypic and genotypic data for both genotyped and non-genotyped individuals can be used to estimate breeding values. Breeding values were estimated from blended pedigree and genomic relationships and progeny canker severity phenotypes for 211 ‘Clapper’ and 154 ‘Graves’ BC_3_F_2_ mothers; from pedigree relationships and progeny phenotypes for 135 ‘Clapper’ and 44 ‘Graves’ BC_3_F_2_ mothers that died prior to genotyping; and from pedigree and genomic relationships alone for 654 ‘Clapper’ and 211 ‘Graves’ BC_3_F_2_ mothers whose progeny had not yet been phenotyped (Fig. 2).

**Figure 2:**
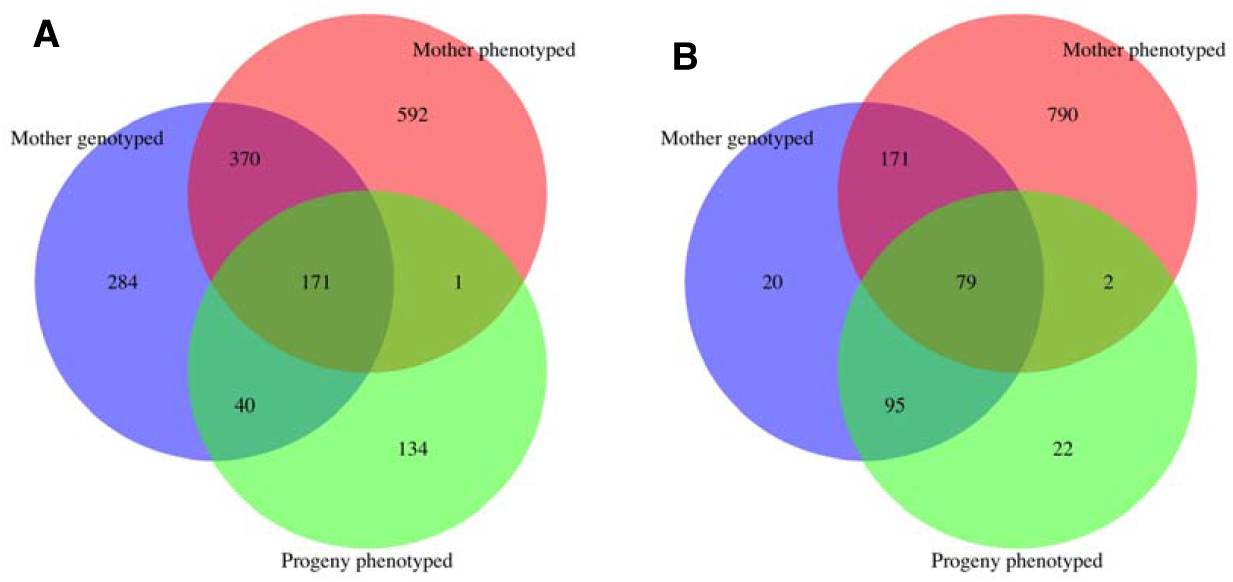
Numbers of BC_3_F_2_ descendants of ‘Clapper’ or ‘Graves’ trees that were phenotyped for five traits indicative of blight tolerance (Mother phenotyped), whose BC_3_F_3_ progeny were phenotyped (Progeny phenotyped), and/or were genotyped for genomic selection (Mother genotyped).

Martini et al. (2018) found that the parameters т and ω that scale the inverse of genomic and pedigree relationships respectively in **H**^−1^, influence predictive ability and inflation of breeding values. We therefore performed single step prediction with H-matrices parameterized with nine pairwise combinations of т and ω involving т = 1, 2, or 3 and ω = 1, 0, or −1 and a tenth combination in which т = ω = 0, which is equivalent to the pedigree relationship matrix. We sought the combination of т and ω that maximized predictive ability while minimizing inflation of breeding values. The inverse of the parameterized H-matrix (hereafter referred to as 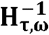) was calculated following Martini et al., (2018):

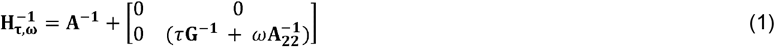

Where **A^−1^** is the inverse of the pedigree relationship matrix, **G**^−1^ is the inverse genomic relationship matrix, and 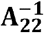 is inverse pedigree relationship matrix among genotyped individuals. Genomic relationships in **G** were estimated following VanRaden (2008):

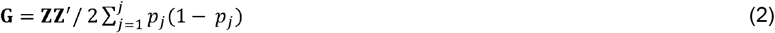

Where **Z** is the centered genotypic matrix and p_j_ are reference allele frequencies for locus 1 through j.

Mixed model analysis with different parameterizations of 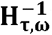 was performed separately for BC_3_F_3_ descendants of ‘Clapper’ and ‘Graves’ populations with the following model in ASReml-R v. 4.1 (Butler et al., 2018):

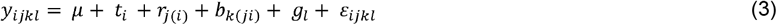

Where y is a vector of canker severity phenotypes for BC_3_F_3_ progeny and μ is the trait mean. The vector t_i_ is composed of the random effects of inoculation years (2011…2016) that were assumed to be independently and normally distributed (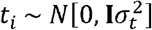, **I** is an identity matrix); **r**_j(i)_ are random effects of complete blocks within the years (2014-2016 trials only) 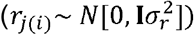; b_k(ij)_ are the random effects of incomplete block within the complete block and year (2014-2016 trials only) 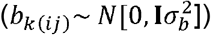; g_l_ are the random additive genetic effects (i.e., the breeding value) of BC_3_F_2_ mothers 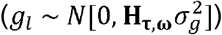; and ε_ijkl_ are the residuals 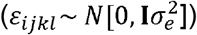. Residuals were approximately normally distributed and no data transformation was performed. The heritability of family mean canker severity 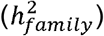 was calculated as:

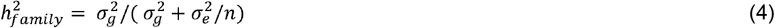

where, n = 13.5 is the mean number BC_3_F_3_ progeny evaluated per BC_3_F_2_ mother tree (Isik et al., 2017).

Genomic predictive ability of breeding values (*r_gĝ_*) for progeny canker severity was estimated with ten-fold cross validation. The cross validation was performed in ASReml-R by randomly subdividing the phenotyped BC_3_F_3_ families into ten subsets and using phenotypic data from 9/10^ths^ of the families to predict breeding values for the remaining 1/10^th^ of the families via 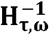. This procedure was repeated for each subset of families to obtain genomic predictions of breeding values for all families. Predictive ability was assessed as the Pearson correlation between the breeding values predicted from genomic and pedigree relationships via 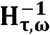 and breeding values estimated with 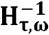 using canker severity data from all phenotyped families. The entire ten-fold cross validation procedure was repeated ten times for each parameterization 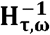 to estimate variation in predictive ability that arises from randomly subdividing the training population into training and prediction subsets.

Inflation of breeding values was estimated from the slope of the regression of adjusted family mean canker severity (y-axis) on the predicted breeding values for progeny canker severity (x-axis) (Martini et al. 2018). Adjusted family means for canker severity were estimated in ASReml-R by treating BC_3_F_2_ mothers as fixed factors and year, block, and incomplete block as random factors as in equation 3. To predict progeny canker severity breeding values, we used the combination of and that maximized predictive ability and where the variation in slope of the regression between adjusted family means and breeding values intersected one among cross validation replicates.

#### Comparing the predictive ability of HBLUP to Bayes C

Predictive ability of the optimized HBLUP procedure was compared to that of Bayes C and prediction from pedigree relationships (ABLUP). Bayes C first estimates the parameter π, which is the proportion of SNPs with non-zero effects and then estimates allelic substitution effects of these SNPs assuming that the effects are normally distributed (Habier et al., 2011). Bayes C was implemented with the R package BGLR (Perez & de los Campos, 2014). Marker effects were estimated over 10,000 iterations of a Gibbs sampler after 2,000 burn-in iterations. To perform ten fold cross validation with Bayes C, allelic substitution effects were estimated on adjusted family mean canker severity for 9/10^th^ of the training population. Genomic estimated breeding values (*ĝ*) for the remaining 1/10^th^ of the population were estimated with:

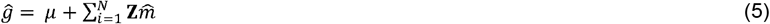

where is **Z** is the centered and imputed genotypic matrix, N is the number of SNPs with minor allele frequency > 0.01, and 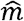 is a vector of allelic substitution effects. To compare predictive ability between methods, predictive ability was estimated as the Pearson correlation between estimated breeding values and adjusted family mean canker severity (*r_yĝ_*).The entire ten-fold cross validation was repeated for ten random partitions of the training population to estimate variation in predictive ability. The same random partitions were used for each method for comparison between methods.

#### Genomic prediction of binary blight phenotypes of BC_3_F_2_ parents

HBLUP analysis of the blight phenotypes of BC_3_F_2_ selection candidates was performed to 1) estimate the heritability and genetic component of these phenotypes and 2) to predict breeding values for genotyped trees age five or less that were too young to reliably express these phenotypes. Breeding values for these traits were predicted for 324 ‘Clapper’ and 115 ‘Graves’ BC_3_F_2_ trees that were age five or younger from genomic or pedigree relationships with 1134 Clapper and 1042 Graves BC_3_F_2_ trees that were phenotyped (Fig. 2). Breeding values and heritability of presence/absence blight phenotypes of individual BC_3_F_2_ trees were estimated with the binomial mixed model:

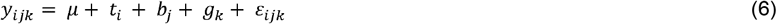

Where y is a binary phenotype (i.e., main stem alive/dead, presence/absence of large cankers, exposed wood, sporulation, or sunken cankers); 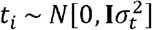 are the random effects of years that the BC_3_F_2_ trees were planted (2002 – 2014); 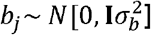 are the random effects of seed orchard block (1…9); 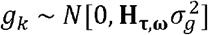 are the random additive genetic effects of individual BC_3_F_2_ tree; and 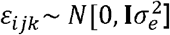 are the residuals. The BC_3_F_2_ phenotypes were coded such that phenotypic classes indicative of blight-tolerance and susceptibility were coded as 1 and 0, respectively (e.g., main stem alive = 1 or dead = 0; large cankers absent = 1 or present = 0; exposed wood absent = 1, present = 0; sporulation absent = 1 or present = 0; and sunken cankers absent = 1, present = 0). Heritability and genomic predictive ability of breeding values were compared for two parameterizations of 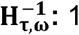) т = ω = 0, which is equivalent to the pedigree relationship matrix and 2) т = ω = 1, which scales **G^−1^** and 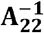 equally. The heritability of blight phenotypes of individual BC_3_F_2_ trees 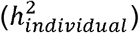 was calculated as:

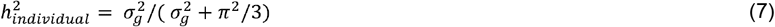

Where *π*^2^/3 is the variance of the standard logistic distribution (Davies et al., 2015). Breeding values for binary blight traits were estimated as probability of having a trait value of 1 given the individual trees’ genotype. This probability was calculated as:

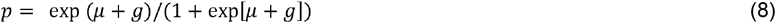

Where μ is the model intercept and g is a vector random genetic effects in units of logit scores (Gezan & Munoz, 2014).

Predictive ability of breeding values for blight phenotypes of individual BC_3_F_2_ trees was assessed using with ten-fold cross validation. Breeding values were predicted for 1/10^th^ of the population from phenotypes and H-matrix relationships with the remaining 9/10ths of the population. Predictive ability was assessed as the Pearson correlation between predicted breeding values (phenotype probabilities) of the genotyped trees when the trees’ phenotypes were left out of the model versus when all trees’ phenotypes were included. The ten-fold cross validation was repeated with ten random partitions of the population for each trait.

### Estimation of blight selection indices and hybrid indices

#### Estimation of selection indices for blight tolerance

Blight selection indices were estimated for all genotyped trees from the sum of from HBLUP breeding values predicted from parent blight phenotypes and progeny canker severity. A selection index called ‘Parent Condition Index’ was created by summing the phenotype probabilities estimated for each of the five blight traits that were phenotyped in the BC_3_F_2_ population. The variance in breeding values for each trait is proportional to the trait’s heritability, thus each trait was weighted in proportion to h^2^_individual_. The breeding values for progeny canker severity were multiplied by −1 to obtain the variable ‘Progeny Blight Tolerance’. Both Parent Condition Index and Progeny Blight tolerance were standardized to mean = 0 and standard deviation = 1 so that they would be equally weighted. The standardized variables were then summed to create the ‘Blight Selection Index.’

#### Estimation of hybrid indices

Hybrid indices were estimated to determine if blight tolerance is correlated with proportion of the backcross trees’ genomes inherited from *C. dentata*. Hybrid indices were estimated for BC_3_F_2_ trees with the R package *introgress* (Gompert & Buerkle, 2010). To generate the required parental data, genotyping-by-sequencing was performed as described above on 56 *C. dentata* individuals and 47 *C. mollissima* individuals. Bioinformatic processing of these data was the same as for the BC_3_F_2_ samples, and after merging data from the pure species and BC_3_F_2_ samples, 27,306 SNPs were retained. The VCF file was converted to STRUCTURE format with PLINK software (http://zzz.bwh.harvard.edu/plink/), and subsequently to *introgress* format using the prepare.data function in *introgress*. Hybrid indices and their confidence limits were then estimated using the est.h function.

## Results

### Accuracy of HBLUP prediction of progeny canker severity

Average predictive ability of breeding values for BC_3_F_3_ progeny canker severity varied from 0.50 to 0.78 for ‘Clapper’ and 0.33 to 0.60 for ‘Graves’ families depending on the scaling parameters т for **G^−1^** and ω for 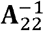 in 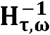 (Table 1). Average predictive ability was lower when predicting from pedigree relationships alone (*r_gĝ_*=0.33 for ‘Clapper’ and *r_gĝ_*=0.41 for ‘Graves’) as compared with most parameterizations of 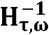 (Table 1). Heritability of BC_3_F_3_ family mean canker severity was maximized for both Clapper 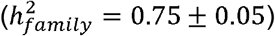 and Graves 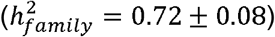 when the scaling parameter for **G^−1^** was maximized (т=3) while the scaling parameter for 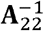 was minimized (ω=−1); however, average inflation of breeding values was also maximized with this parameterization of 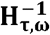 (Table 1). The combination of scaling parameters that maximized predictive ability with inflation values that intersected one across cross validation replicates was т = 3 and ω = 1 (Table 1). Many of the genotyped BC_3_F_2_ trees were more closely related than expected from pedigree relationships (Fig. A3), which may explain the increased predictive ability when pedigree and genomic relationships were blended in 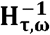.

**Table 1:** The effect of different parameterizations of the 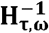 inverse relationship matrix on family mean heritability (h^2^_family_), predictive ability of breeding values (*r_gĝ_*), and inflation of breeding values for canker severity among BC_3_F_2_ descendants of the ‘Clapper’ and ‘Graves’ trees. The parameters т and ω scale pedigree and genomic relationships, respectively, among genotyped individuals.

| $\tau$ | $\omega$ | $h^2_{\text{family}} \pm \text{SE}$ | | Predictive ability ($r_{g\hat{g}}$) | | | | | | Inflation of breeding values | | | | | |
| --- | --- | --- | --- | --- | --- | --- | --- | --- | --- | --- | --- | --- | --- | --- | --- |
|  |  | Clapper | Graves | Clapper |  |  | Graves |  |  | Clapper |  |  | Graves |  |  |
|  |  |  |  | Max | Avg | Min | Max | Avg | Min | Max | Avg | Min | Max | Avg | Min |
| 3 | 1 | 0.67 ± 0.06 | 0.59 ± 0.09 | 0.79 | 0.78 | 0.75 | 0.63 | 0.60 | 0.58 | 1.16 | 1.09 | 0.98 | 1.60 | 1.27 | 0.94 |
| 2 | 1 | 0.64 ± 0.06 | 0.50 ± 0.10 | 0.76 | 0.74 | 0.72 | 0.62 | 0.60 | 0.57 | 1.10 | 1.03 | 0.92 | 1.50 | 1.19 | 0.88 |
| 1 | 1 | 0.54 ± 0.07 | 0.34 ± 0.09 | 0.71 | 0.70 | 0.67 | 0.62 | 0.60 | 0.58 | 1.03 | 0.95 | 0.84 | 1.43 | 1.14 | 0.84 |
| 3 | 0 | 0.73 ± 0.05 | 0.68 ± 0.08 | 0.70 | 0.68 | 0.65 | 0.46 | 0.43 | 0.40 | 1.53 | 1.39 | 1.28 | 2.33 | 1.69 | 1.05 |
| 2 | 0 | 0.71 ± 0.05 | 0.65 ± 0.09 | 0.65 | 0.63 | 0.60 | 0.43 | 0.40 | 0.36 | 1.48 | 1.33 | 1.21 | 2.26 | 1.63 | 0.97 |
| 1 | 0 | 0.68 ± 0.05 | 0.58 ± 0.09 | 0.56 | 0.55 | 0.51 | 0.38 | 0.35 | 0.31 | 1.39 | 1.21 | 1.06 | 2.13 | 1.54 | 0.87 |
| 3 | -1 | 0.75 ± 0.05 | 0.72 ± 0.08 | 0.65 | 0.63 | 0.60 | 0.42 | 0.39 | 0.34 | 1.66 | 1.48 | 1.32 | 2.71 | 1.92 | 1.11 |
| 2 | -1 | 0.74 ± 0.05 | 0.69 ± 0.08 | 0.60 | 0.58 | 0.55 | 0.39 | 0.36 | 0.31 | 1.57 | 1.37 | 1.19 | 2.58 | 1.84 | 1.03 |
| 1 | -1 | 0.72 ± 0.05 | 0.65 ± 0.09 | 0.52 | 0.50 | 0.46 | 0.36 | 0.33 | 0.28 | 1.37 | 1.15 | 0.95 | 2.31 | 1.69 | 0.92 |
| 0 | 0 | 0.56 ± 0.06 | 0.35 ± 0.10 | 0.35 | 0.33 | 0.29 | 0.44 | 0.41 | 0.38 | 0.63 | 0.51 | 0.40 | 1.45 | 1.13 | 0.73 |

Note: The pedigree relationship matrix is equivalent to $\tau = \omega = 0$ .

### Prediction of progeny canker severity using Bayes C

There was no gain in predictive ability of BC_3_F_3_ family mean canker severity using Bayes C, which sets a proportion of the marker effects to zero, as compared with HBLUP, which incorporates all markers into the prediction (Table 2). This result suggests that an infinitesimal model is as accurate as a major gene model for predicting blight tolerance. Predictive ability of family mean canker severity averaged across the HBLUP and Bayes C methods was greater for the ‘Clapper’ population (*r_yĝ_* = 0.256) relative to the ‘Graves’ population (*r_yĝ_* = 0.155). Higher predictive abilities in ‘Clapper’ versus ‘Graves’ populations, respectively, may be attributed to the larger training population in ‘Clapper’ (211 v. 154 genotyped BC_3_F_2_ mothers; Fig. 2) and higher heritability of BC_3_F_3_ family mean canker severity in ‘Clapper’ (0.67 ± 0.06 v. 0.59 ± 0.09) (Table 1). Both and HBLUP and Bayes C methods were more accurate than prediction from the pedigree (ABLUP) (Table 2).

**Table 2:** Comparison of the predictive ability of different genomic selection methods. Predictive ability was assessed with ten-fold cross as the correlation between the predicted and observed BC_3_F_3_ family mean canker severity (*r_yĝ_*). To estimate variation in predictive ability, the cross validation repeated ten times with different random partitions of the training population. In the Bayes B and Bayes C methods, the proportion of markers with non-zero effects varied between cross validation replicates.

|  | Clapper |  |  |  |  |  | Graves |  |  |  |  |  |
| --- | --- | --- | --- | --- | --- | --- | --- | --- | --- | --- | --- | --- |
| | Proportion of markers included | | | Predictive ability ($r_{y\hat{g}}$) | | | Proportion of markers included | | | Predictive ability ($r_{y\hat{g}}$) | | |
|  | Max | Avg | Min | Max | Avg | Min | Max | Avg | Min | Max | Avg | Min |
| HBLUP | 1 | 1 | 1 | 0.30 | 0.28 | 0.25 | 1 | 1 | 1 | 0.19 | 0.15 | 0.11 |
| ABLUP | 0 | 0 | 0 | 0.24 | 0.21 | 0.18 | 0 | 0 | 0 | 0.10 | 0.06 | 0.02 |
| Bayes C | 0.54 | 0.27 | 0.14 | 0.28 | 0.25 | 0.20 | 0.49 | 0.31 | 0.16 | 0.22 | 0.16 | 0.10 |

### Heritability and predictive ability of blight phenotypes of BC_3_F_2_ parents

Blight tolerance phenotypes of BC_3_F_2_ trees (age 5 to 17) were weakly heritable, with h^2^_individual_ values varying from 0 to 0.25 depending on the trait (Table 3). Heritabilities estimated with the H-matrix were similar to those estimated with the pedigree. On average, trees with an observed phenotype indicative of blight tolerance (i.e., main stem alive) also had a greater probability of expressing a blight tolerance phenotype given the tree’s genotype (Fig. 3). However, due to the low heritability of the blight phenotypes of BC_3_F_2_ trees, there was overlap in the distributions phenotype probabilities between observed resistant versus susceptible phenotype classes. In the ‘Graves’ population, there was no difference in phenotypic probabilities for the presence/absence of large cankers and presence/absence of sporulation, suggesting these traits were not informative (Fig. 3). Averaged across traits, predictive ability was 1.3 times greater in the ‘Clapper’ population and 2.5 times greater in the ‘Graves’ population when using the H-matrix as compared with prediction from the pedigree (Table 3).

**Figure 3:**
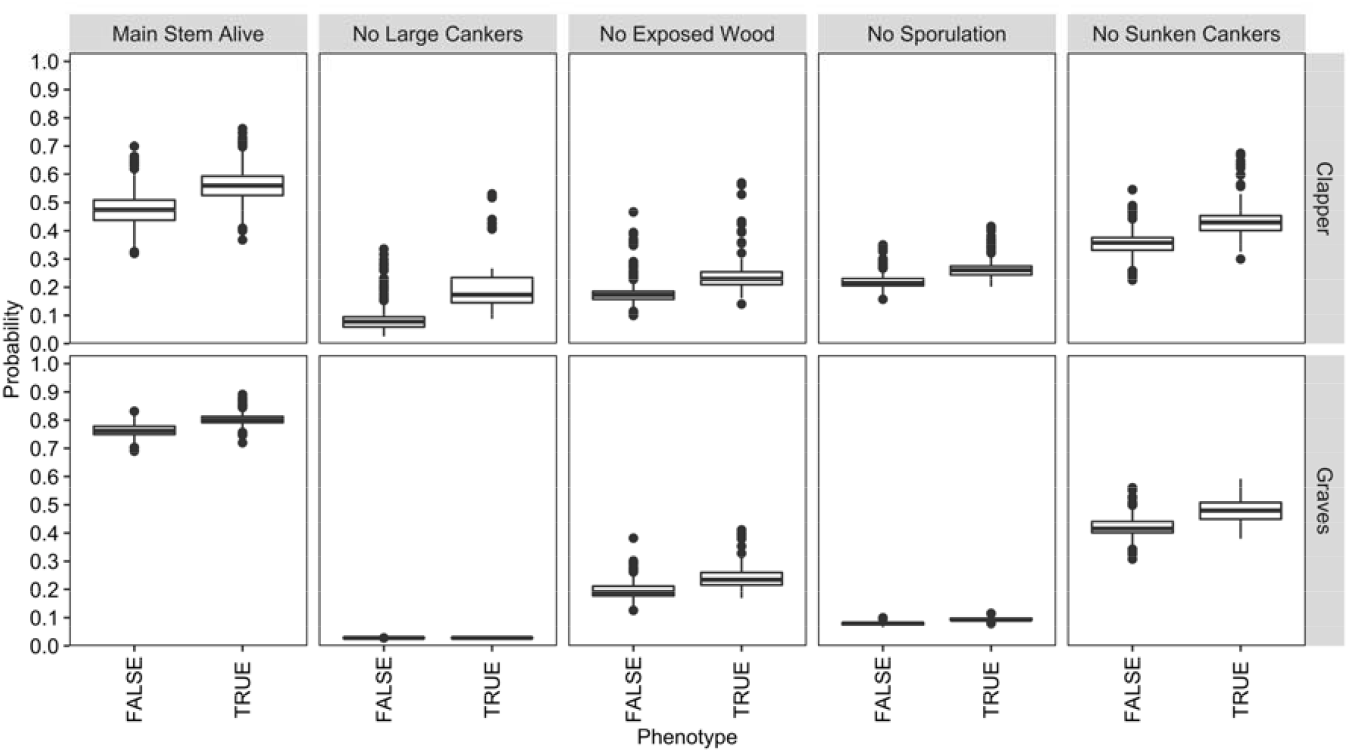
Probabilities that BC_3_F_2_ trees will have a phenotype indicative of blight tolerance given trees’ genotypes versus trees’ observed phenotypes for ‘Clapper’ and ‘Graves’ BC_3_F_2_ trees.

**Table 3:** Comparison of the heritability and predictive ability of binary blight phenotypes of BC_3_F_2_ trees. Predictive ability was assessed with ten-fold cross validation as the correlation between phenotype probabilities when the phenotype was observed v. left out. The ten-fold cross validation was repeated ten times with different random partitions of the training population to estimate variation in predictive ability.

|  | N TRUE | N FALSE | Pedigree<br>h<sup>2</sup> ± SE | H-matrix<br>h<sup>2</sup> ± SE | Pedigree<br>predictive ability |  |  | H-matrix<br>predictive ability |  |  |
| --- | --- | --- | --- | --- | --- | --- | --- | --- | --- | --- |
|  |  |  |  |  | Max | Avg | Min | Max | Avg | Min |
| *Clapper* |  |  |  |  |  |  |  |  |  |  |
| Main stem alive | 645 | 489 | 0.07 ± 0.04 | 0.08 ± 0.04 | 0.78 | 0.75 | 0.67 | 0.93 | 0.91 | 0.89 |
| No large cankers | 55 | 577 | 0.20 ± 0.10 | 0.25 ± 0.09 | 0.78 | 0.74 | 0.68 | 0.91 | 0.88 | 0.83 |
| No sporulation | 151 | 483 | 0.03 ± 0.07 | 0.07 ± 0.06 | 0.52 | 0.44 | 0.27 | 0.90 | 0.86 | 0.81 |
| No exposed wood | 109 | 519 | 0.11 ± 0.08 | 0.12 ± 0.06 | 0.80 | 0.70 | 0.60 | 0.94 | 0.92 | 0.88 |
| No sunken cankers | 245 | 388 | 0.08 ± 0.06 | 0.09 ± 0.05 | 0.76 | 0.72 | 0.67 | 0.92 | 0.90 | 0.89 |
| *Graves* |  |  |  |  |  |  |  |  |  |  |
| Main stem alive | 872 | 170 | 0.08 ± 0.06 | 0.06 ± 0.06 | 0.42 | 0.37 | 0.34 | 0.84 | 0.80 | 0.73 |
| No large cankers | 28 | 843 | 0.05 ± 0.24 | 0 | 0.32 | 0.26 | 0.17 | NA | NA | NA |
| No sporulation | 70 | 802 | 0.04 ± 0.12 | 0.05 ± 0.11 | 0.28 | 0.21 | 0.13 | 0.60 | 0.55 | 0.50 |
| No exposed wood | 191 | 681 | 0.04 ± 0.05 | 0.07 ± 0.05 | 0.36 | 0.31 | 0.25 | 0.95 | 0.94 | 0.92 |
| No sunken cankers | 412 | 457 | 0.07 ± 0.04 | 0.07 ± 0.05 | 0.52 | 0.46 | 0.40 | 0.89 | 0.87 | 0.86 |

Note: predictive ability using the H-matrix was not assessed for presence/absence of large cankers in the 'Graves' population because the trait heritability was zero.

### Estimation of hybrid indices

Hybrid indices varied from 0.9996 (nearly 100% *C. dentata)* to 0.4171 (58% *C. mollissima)* for 865 BC_3_F_2_ descendants of ‘Clapper’, and from 0.9996 to 0.3463 for 365 BC_3_F_2_ descendants of ‘Graves’ (Fig. 4). There were 24 ‘Clapper’ and 10 ‘Graves BC_3_F_2_ trees with hybrid indices less than or equal 0.55. These trees were inferred to be ‘pseudo-F_1_’ progeny of BC_3_ mother trees that were pollinated by *C. mollissima* trees on the same property. The average hybrid index of ‘Clapper’ and ‘Graves’ BC_3_F_2_ trees, excluding pseudo-F_1_s, was 0.8943.

**Figure 4:**
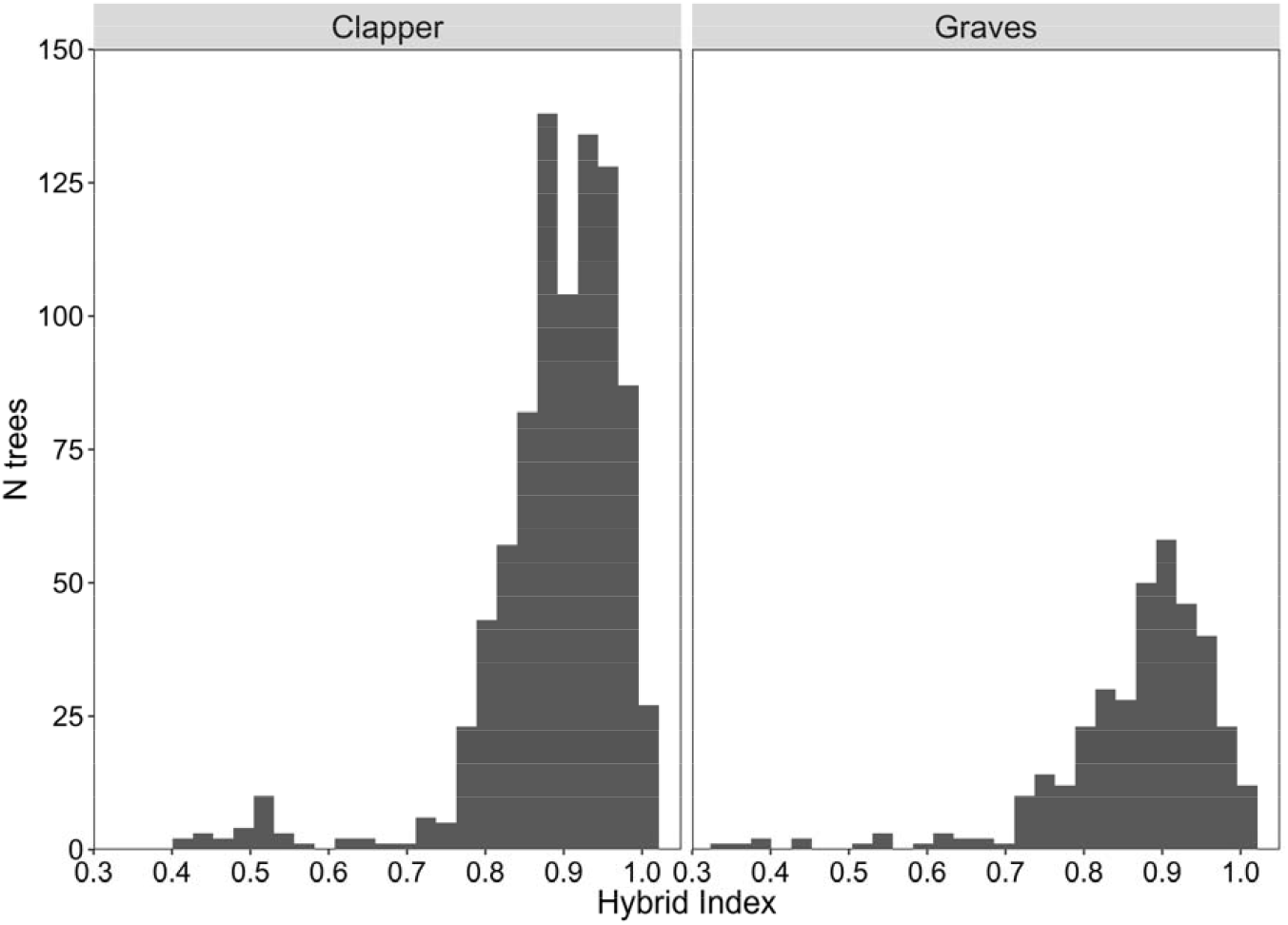
Distribution of hybrid index values for BC_3_F_2_ descendants of ‘Clapper’ and ‘Graves’. Hybrid index values indicate the proportion of hybrid genomes inherited from *C. dentata* v. *C. mollissima* (1 = 100% *C. dentata*).

### Comparison of different selection scenarios

Blight Selection Indices for genotyped trees were obtained by summing the Parent Condition Index and Progeny Blight Tolerance (see Materials & Methods). These selection candidates from the ‘Clapper’ and ‘Graves’ populations were planted in 161 and 116 seed orchard plots, respectively. We considered three selection scenarios: 1. Select one tree within each seed orchard plot with the largest Blight Selection Index. 2. Select an equal number of trees, but select trees with the largest Blight Selection Index regardless of seed orchard plot. 3. Select the same number of trees, but select a maximum of three trees per plot with the largest Blight Selection Indices. The pseudo-F_1_ trees were excluded from consideration for selection; however, Blight Selection Indices of the selected trees were compared to that of the pseudo-F_1_s.

For both the ‘Clapper’ and ‘Graves’ populations, all selections scenarios were predicted to increase the mean Blight Selection Index. However, selected trees were, on average, significantly less blight-tolerant than pseudo-F_1_s (Fig. 5). The average Blight Selection Index of selected BC_3_F_2_ trees was significantly greater when selecting trees with the maximum Blight Selection Index (Scenario 2) or selecting up to three trees per plot (Scenario 3) as compared with selecting one tree per plot (Scenario 1).

**Figure 5:**
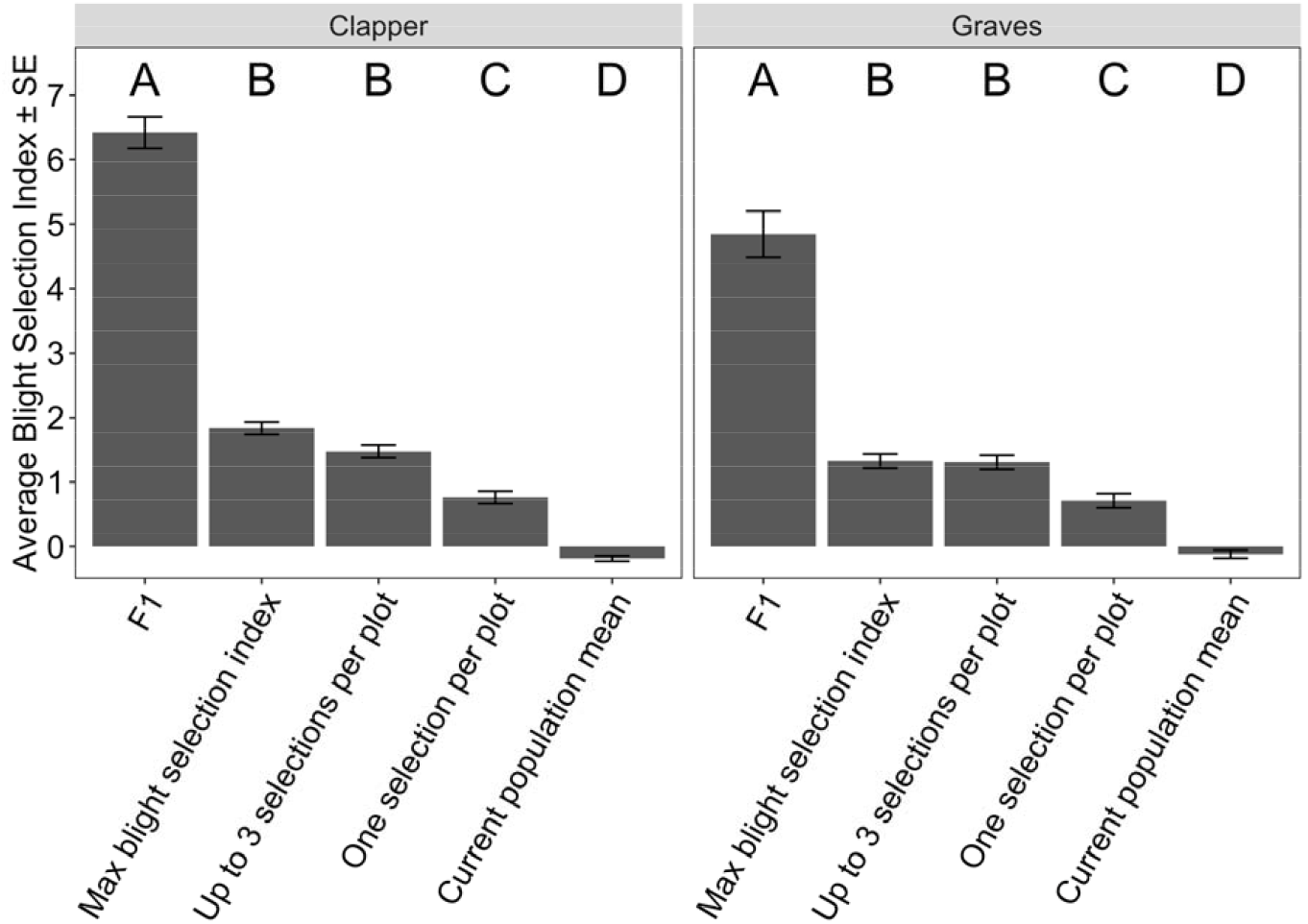
Comparison of average Blight Selection Indices for selected ‘Clapper’ and ‘Graves’ BC_3_F_2_ trees under different selection scenarios. Selection scenarios included making one selection per 150 half sibs planted in each seed orchard subplot (one selection per plot); selecting an equal number of trees with the maximum Blight Selection Index (max blight selection index); and making up to three selections per plot (up to three selections per plot). The average Blight Selection Index for the selected BC_3_F_2_ trees was compared to that of the current population and pseudo-F_1_ trees (i.e., progeny of BC_3_ trees outcrossed to *C. mollissima*). Letters above the bars indicate the significance of differences in average Blight Selection Index (Tukey test, P < 0.05).

The tradeoff when relaxing the constraint of selecting one tree per plot was a reduction of the number of *C. dentata* backcross lineages represented among the selections. For example, the ‘Clapper’ BC_3_F_2_ selection candidates had 41 and 28 *C. dentata* grandparents and great-grandparents in their maternal line. By selecting 160 trees with the maximum Blight Selection Indices regardless of plot (Scenario 2), selections included descendants from 31 *C. dentata* grandparents and 24 great-grandparents. By selecting a maximum of three trees per plot, selections included descendants of 33 grandparents and 25 great-grandparents. We decided to proceed with up to three selections per plot because this scenario resulted in selections with a similar average Blight Selection Indices as Scenario 2 (Fig. 5), but retained a larger proportion of the maternal *C. dentata* lineages.

For both the ‘Clapper’ and ‘Graves’ populations, blight-tolerance as assessed with the Parent Condition Index, Progeny Blight Tolerance, and Blight Selection Index was negatively correlated with the proportion of alleles inherited from *C. dentata* (Fig. 6). These negative correlations were observed when genomic prediction models were developed with and without including pseudo-F_1_s in the training population, suggesting that the pseudo-F_1_s are not driving this result (not shown). Selected BC_3_F_2_ trees were estimated to have inherited an average (max, min) of 83% (99%, 61%) of their genome from *C. dentata*. Parent Condition Index was positively correlated with Progeny Blight Tolerance (Fig. 7). A total of 121 of 161 ‘Clapper’ and 70 of 116 ‘Graves’ selections had above average Parent Condition Index and above average Progeny Blight Tolerance (Fig. 7). A representative BC_3_F_2_ selection, a pseudo-F_1_, and a pure *C. dentata* are pictured in Fig. A4.

**Figure 6:**
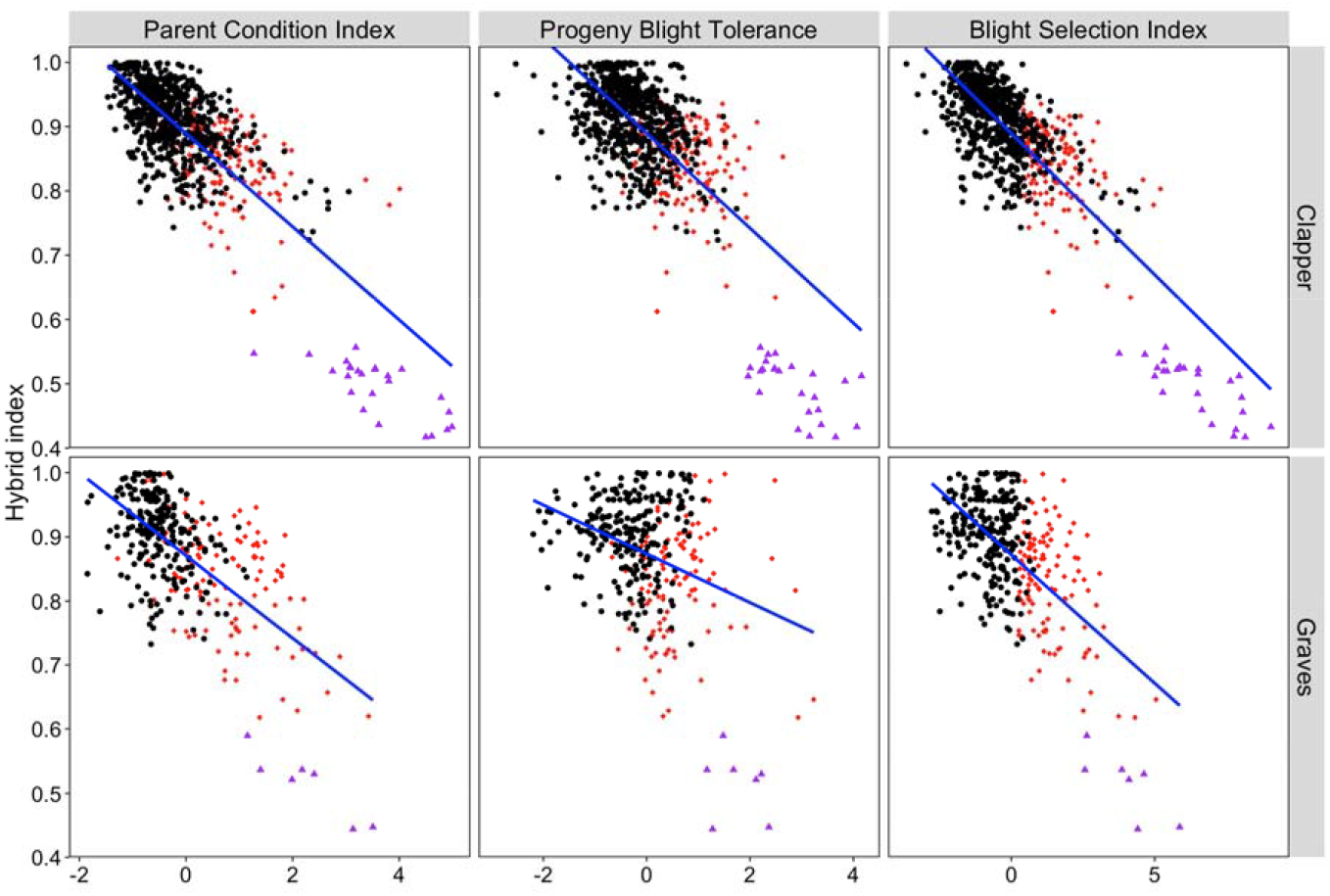
Proportion of Clapper and Graves BC_3_F_2_ genomes inherited from *C. dentata* (hybrid index) versus blight-tolerance. Blight-tolerance was assessed via the Parent Condition Index (a sum of phenotype probabilities for in late-developing blight trait on BC_3_F_2_ stems), Progeny Blight Tolerance (breeding values for average progeny canker severity, reversed in scale), and Blight Selection Index (Parent Condition Index + Progeny Blight Tolerance). Red triangles are BC_3_F_2_ selections (up to three selections per 150-tree subplot), purple diamonds are the pseudo-F_1_ progeny of BC_3_ trees outcrossed to *C. mollissima*, and black dots are inferior trees to cull. Blue lines are the least squares regressions between hybrid index and blight-tolerance traits.

**Figure 7:**
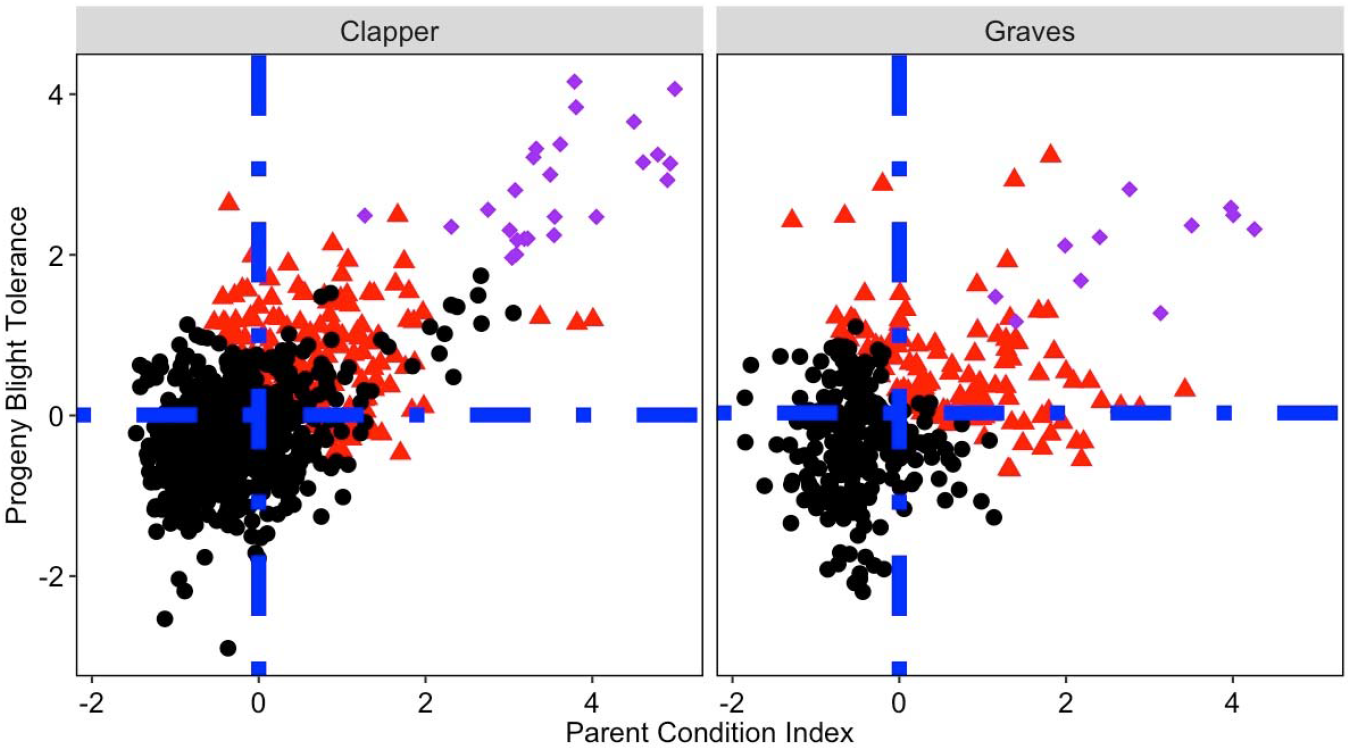
Relationship between Parent Condition Index and Progeny Blight Tolerance for ‘Clapper’ and ‘Graves’ populations. Red triangles are BC_3_F_2_ selections with up to three selections per seed orchard plot, purple diamonds are pseudo-F_1_ progeny of BC_3_ trees outcrossed to *C. mollissima*, and black dots are inferior trees to cull. Blue dashed lines are the population means for Parent Condition Index and Progeny Blight Tolerance.

## Discussion

### Outlook for genomic selection of the most blight-tolerant trees in BC_3_F_2_ seed orchards

Our first aim in this study was to optimize genomic selection to increase the speed and accuracy of making final selections for blight-tolerance in American chestnut BC_3_F_2_ seed orchards. We integrated two generations (BC_3_F_2_ and BC_3_F_3_) of blight-tolerance phenotypes with genotyping-by-sequencing of BC_3_F_2_ trees to select the most blight-tolerant trees. We were successful insofar as the accuracy of genomic prediction of both BC_3_F_2_ blight tolerance phenotype probabilities and breeding values for BC_3_F_3_ progeny canker severity was more accurate than prediction from pedigree relationships. Furthermore, all selection scenarios were predicted to increase the average blight tolerance relative to the current population mean.

We plan to finish selection in the Meadowview seed orchards in the next few years with additional progeny testing and genomic selection. An additional 184 ‘Clapper’ and 216 ‘Graves’ BC_3_F_3_ families will be inoculated in field trials in 2019, 2020, and 2021. Furthermore, genotyping of approximately 1,000 additional ‘Clapper’ and ‘Graves’ BC_3_F_2_ trees is currently ongoing. We anticipate that the accuracy of genomic selection will increase by expanding the training population as has been predicted from simulation studies (Grattipaglia & Resende, 2011) and observed for other species and traits (Asoro et al. 2011; Zhang et al., 2017).

### Evaluating the major gene hypothesis for blight tolerance

Our second aim was to use genomic selection and hybrid index analysis to evaluate the hypothesis of major gene inheritance of blight tolerance. Two observations support the alternative hypothesis that blight tolerance is inherited as a polygenic trait. First, we observed a tradeoff between blight tolerance and the proportion of BC_3_F_2_ trees’ genomes inherited from *C. dentata*. Second, HBLUP, which assumes an infinitesimal model of inheritance, was just as accurate at predicting progeny canker severity as Bayes C, which includes only the markers with largest effects in the prediction model. Previous QTL mapping studies of blight tolerance were conducted in a small *C. dentata* x *C. mollissima* F_2_ family (<100 full sib progeny) (Kubisiak et al., 1997; Kuibisiak et al., 2013); therefore it is likely that the effects of individual loci were inflated and these studies were underpowered to comprehensively detect all loci associated with blight-tolerance (Beavis, 1994; Slate, 2013). Regardless of the number of loci underlying blight-tolerance, the low heritabilities (h^2^ < 0.25) of blight-tolerance phenotypes of suggests that some alleles for blight-tolerance have been lost in some backcross generations and lines as a result of low-accuracy phenotypic selection.

### Revised projections of average blight-tolerance after selection at BC_3_F_2_

Steiner et al. (2017) predicted final selection in BC_3_F_2_ seed orchards would result in a BC_3_F_3_ population with an average blight-tolerance similar to *C. mollissima* x *C. dentata* F_1_ hybrids. We observed that average blight tolerance of BC_3_F_2_ selections that inherited approximately 90% of their genome from *C. dentata* was less than that of pseudo-F_1_ trees, which inherited approximately 50% of their genome from *C. dentata*. Previous studies have found that BC_3_F_3_ progeny from partially selected seed orchards have improved blight-tolerance relative to *C. dentata* in orchard and greenhouse trials (Steiner et al., 2017; Westbrook & Jarrett, 2018), Therefore, we predict that the average blight-tolerance of BC_3_F_3_ progeny from fully selected BC_3_F_2_ seed orchards will be between that of F_1_ hybrids and *C. dentata*.

### Where does breeding for American chestnut restoration go from here?

#### Restoration trials

It is not known what combination of blight-tolerance and *C. dentata* inheritance will be sufficient for American chestnut restoration. The American chestnut Foundation has planted field trials composed of BC_3_F_3_ progeny from Meadowview seed orchards at over 35 sites across the eastern U.S. (Fig. 1). Many of these trials are between five and ten years old: too young to reliably assess for blight tolerance following natural infection by *C. parasitica*. Encouragingly, in the oldest field trials, blight incidence and severity on eight-year old BC_3_F_3_ trees was lower than on pure American chestnut and similar to Chinese chestnut (Clark et al., 2019). Once selection is complete in seed orchards, TACF intends to plant additional restoration trials composed of the most blight tolerant BC_3_F_3_ families planted on sites most suitable for growing American chestnut.

The influence of environmental factors such as competition, climate, and soil on blight-tolerance will be estimated via replication of BC_3_F_3_ families across sites and varying silvicultural treatments within sites (TACF, 2012).

#### Selection for blight tolerance and timber-type form in earlier backcross generations

The American Chestnut Foundation is currently generating and selecting *C. dentata* backcross progeny from ten additional *C. mollissima* sources of blight-tolerance (Steiner et al., 2017; Westbrook et al., 2018). Based on the finding of a tradeoff between blight-tolerance and *C. dentata* inheritance, we will advance these additional sources only to the BC_1_ or BC_2_ generations rather than BC_3_ before intercrossing the selections. Backcross trees will be selected for blight-tolerance not only with phenotypic selection, but also by inoculating progeny derived from controlled pollinations of these trees to ensure that selection is accurate.

While BC_1_ and BC_2_ selections are expected to be more blight tolerant than selections from later backcross generations, the earlier backcross selections are expected to inherit other traits from *C. mollissima* that may be undesirable for forest restoration. Compared with American chestnut, Chinese chestnuts growing in North America generally have lower height growth (Diller & Clapper, 1969; Sclarbaum et al. 1998; Thomas-Van Gundy, 2016), greater stem branching (Clark et al., 2012), lower maximum photosynthetic rates (Knapp et al., 2014), lower cold tolerance (Gurney et al., 2011; Saielli et al., 2012), and differential colonization of roots by mycorrhizae and other fungi (Reazin et al., 2019). After selecting for blight tolerance, we will perform additional selection for timber-type form and overall proportion of backcross trees’ genomes inherited from *C. dentata*, which will necessitate screening large populations segregating for these traits.

#### Incorporating transgenic blight-tolerance

Lower than expected blight-tolerance within BC_3_F_3_ populations highlights potential advantages of using transgenic American chestnut trees for restoration. Transgenic *C. dentata* founder lines that constitutively overexpress an oxalate oxidase (OxO) gene from wheat have high levels of blight-tolerance in seedling trials (Newhouse et al., 2014; Powell et al., 2019). Progeny from transgenic x wild-type crosses essentially inherit 100% of their genome from *C. denata*. The inheritance of OxO, which is expected in approximately 50% of the progeny, can be detected inexpensively with an enzymatic assay or with PCR (Zhang et al., 2013). Federal regulatory review in the United States is ongoing to release transgenic American chestnut founder trees for breeding and restoration trials outside of a few confined, permitted field trials. If federal regulatory approval is granted, TACF plans to outcross transgenic founder clone(s) to wild-type trees over five generations to increase the effective population size to > 500 and to maximize genome inheritance from wild-type trees with marker-assisted introgression (Westbrook et al., 2019). Transgenic trees may also be crossed with backcross trees to potentially enhance blight-tolerance. Public acceptance of transgenic American chestnut trees for restoration is mixed (Delbourne et al., 2018) and the long-term blight-tolerance of transgenic trees in forest conditions is not currently known. Therefore, it is prudent to continue traditional breeding approaches to introgress blight-tolerance from Asian *Castanea* species into *C. denata* separately from breeding with transgenic trees.

## Conclusions and future directions

In developing genomic prediction models and estimating hybrid indices for BC_3_F_2_ American chestnuts, we discovered a tradeoff between blight-tolerance and proportion of the genome inherited from *C. dentata*. Results suggest that genetic architecture underlying the inheritance of blight-tolerance is more complex than previously assumed. A chromosome-scale genome assembly for *Castanea dentata* is forthcoming, which will be combined with genotyping of thousands of backcross individuals to enable mapping the inheritance of *C. mollissima* haplotypes and discovery of genomic regions associated with variation in blight-tolerance.

### Data archiving statement

Demultiplexed and quality-trimmed sequence reads per sample have been uploaded to the NCBI Sequence Read Archive (SRA) under bioproject accessions PRJNA507748 and PRJNA507747. The blight phenotypes and a VCF file containing the filtered and imputed SNPs can be accessed on Dryad. Contact Jeremy Schmutz for access to the latest assembly of the *Castanea dentata* genome sequence under the Ft. Lauderdale agreement. Once the annotation is finalized, the genome will be publicly available at the Phytozome, comparative plant genomics portal.

## Acknowledgements

We would like to thank the donors and volunteers with The American Chestnut Foundation who have supported the breeding effort for the past 35 years. We also thank Advanced Research Computing at Virginia Tech for providing computational resources and technical support related to the analyses described here. Initial funding for proof-of-concept for genomic selection was provided by the U.S. Endowment for Forestry and Communities and the United States Forest Service. Subsequent funding to develop genomic prediction models was provided by USDA-NIFA Project 2016-67013-24581 and McIntire-Stennis Project 1005394. Funding for genotyping to predict resistance of remaining trees in Meadowview seed orchards was provided by the Allegheny Foundation and an anonymous foundation. The Colcom Foundation provided funding to generate the draft reference genome for American chestnut.

---

**Figure A1:**
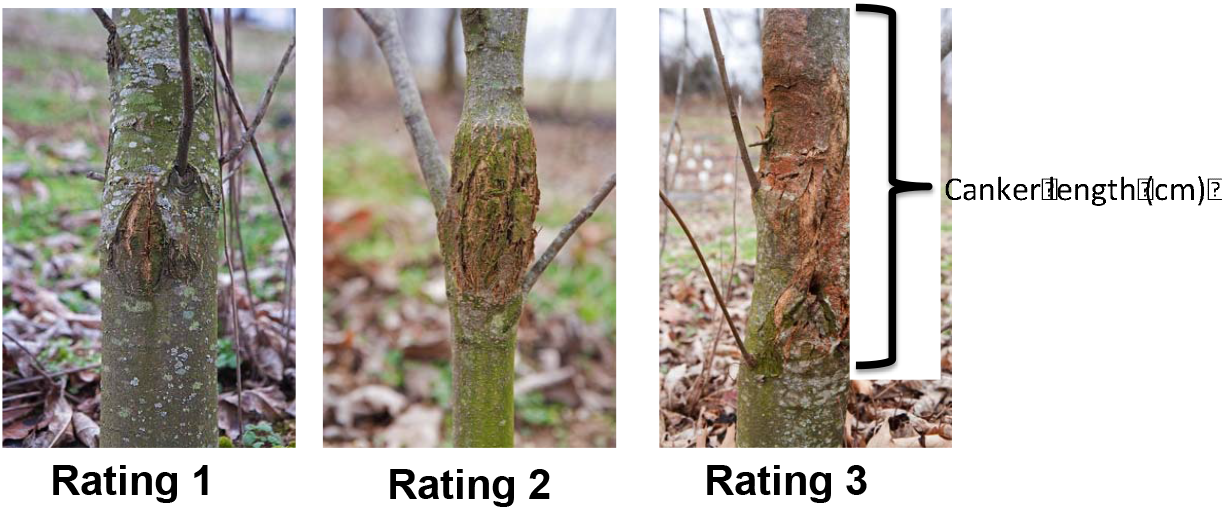
Pictures of subjective canker ratings and canker lengths obtained when phenotyping American chestnut BC_3_F_3_ trees for blight-tolerance.

**Figure A2:**
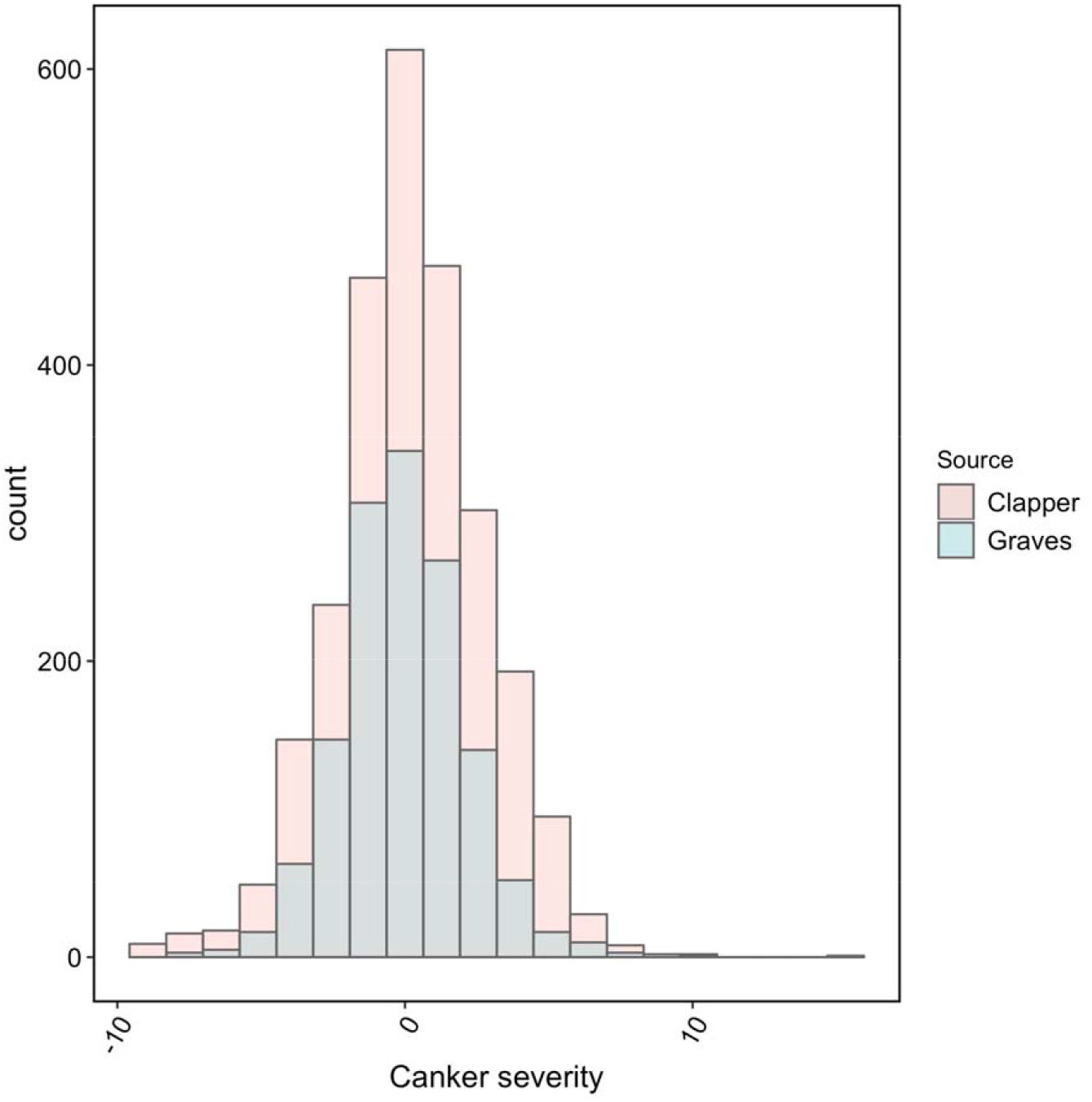
Distribution of canker severity values for BC_3_F_3_ descendants of ‘Clapper’ and ‘Graves.’

**Figure A3:**
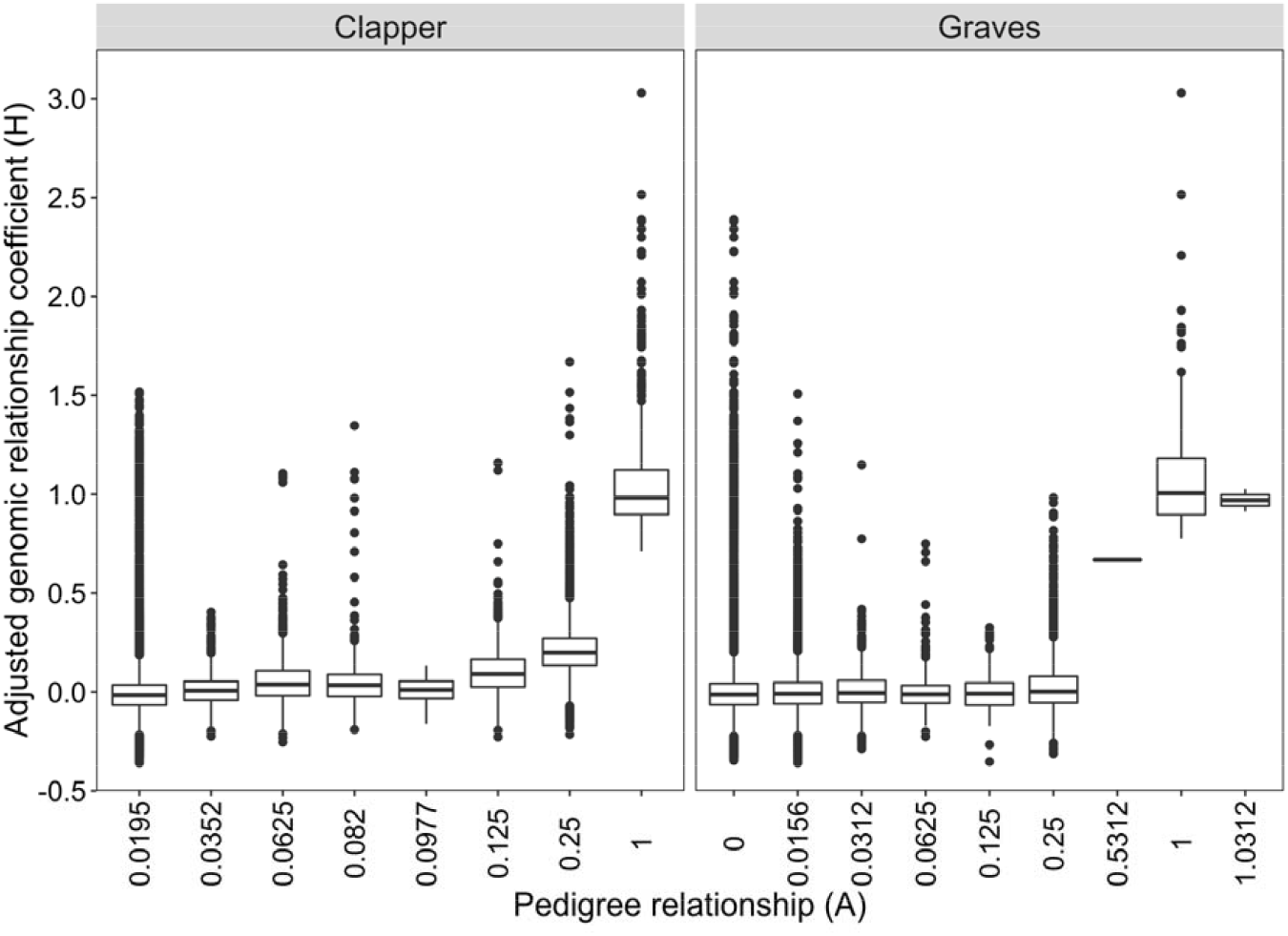
Comparison of pedigree and adjusted genomic relationship coefficients for BC_3_F_2_ descendants of ‘Clapper’ and ‘Graves.’ Genomic relationship coefficients were centered on their expected pedigree values with pedigree and genomic relationships scaled equally (т = ω = 1).

**Figure A4:**
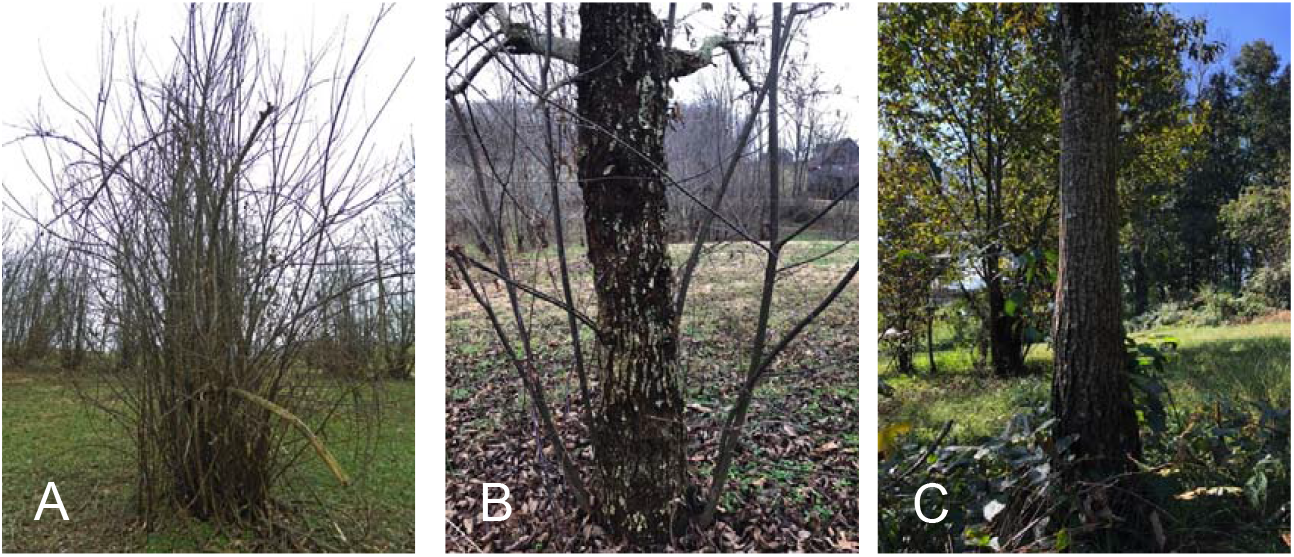
A pure *Castanea dentata* (A) as compared with a *C. dentata* BC_3_F_2_ hybrid selections (B) and a pseudo-F_1_ (*C. dentata* BC_3_ x *C. mollissima*) (C).

